# Episodic evolution of a eukaryotic *NADK* repertoire of ancient provenance

**DOI:** 10.1101/507913

**Authors:** Oliver Vickman, Albert Erives

## Abstract

NAD kinase (NADK) is the sole enzyme that phosphorylates nicotinamide adenine dinucleotide (NAD+/NADH) into NADP+/NADPH, which provides the chemical reducing power in anabolic (biosynthetic) pathways. While prokaryotes typically encode a single NADK, eukaryotes encode multiple NADKs. How these different *NADK* genes are all related to each other and those of prokaryotes is not known. Here we conduct phylogenetic analysis of *NADK* genes and identify major clade-defining patterns of *NADK* evolution. First, almost all eukaryotic *NADK* genes belong to one of two ancient eukaryotic sister clades corresponding to cytosolic (“cyto”) and mitochondrial (“mito”) clades. Secondly, we find that the cyto-clade *NADK* gene is duplicated in connection with loss of the mito-clade NADK gene in several eukaryotic clades or with acquisition of plastids in Archaeplastida. Thirdly, we find that horizontal gene transfers from proteobacteria have replaced mitochondrial *NADK* genes in only a few rare cases. Last, we find that the eukaryotic cyto and mito paralogs are unrelated to independent duplications that occurred in sporulating bacteria, once in mycelial Actinobacteria and once in aerobic endospore-forming Firmicutes. Altogether these findings show that the eukaryotic *NADK* gene repertoire is ancient and evolves episodically with major evolutionary transitions.

**Author Summary:** Metabolic enzymes are central to all living organisms and key to understanding the evolution of life on Earth. One important component of all metabolic pathways is the participation of chemically-reducing cofactors such as the redox system of NAD+ and NADH molecules. All living cells also possess a system for shifting their intracellular pool of NAD+ and NADH into NADP+ and NADPH via phosphorylation by the NAD kinase (NADK). Phosphorylation by NADK allows cells to regulate the use of their main chemical redox system for anabolic growth and the synthesis of basic cellular components. In our study we provide the first comprehensive analysis of the NAD kinase family. We find that eukaryotes possess two ancient NADK gene families of unknown provenance and also find unrelated bacterial families that also possess dual NADK genes. We also find that while NADK evolution has been mostly stable through the eons, this family has evolved substantially in association with the evolution of specific clades of organisms. Altogether these results provide a metabolic picture of the tree of life and its evolution.

## Introduction

Nicotinamide adenine dinucleotide (NAD) is a dinucleotide cofactor essential to all living cells (1–7). NAD is composed of one nucleotide featuring the pyrimidine base nicotinamide (N) and another nucleotide featuring the purine base adenine (A). The nicotinamide base of NAD is an electron carrier central to NAD’s function. In the oxidized form, NAD is written as “NAD+” to emphasize it is in the unreduced state. The reduced form occurs after acquisition of two electrons and 1 proton (+ 2e^-^ + H^+^) and is written as “NADH”. NADH is a strong, biologically-relevant, electron donor such that a 1- to-1 mixture of NADH and NAD+ has a redox potential of −320 mV (5). We can diagram the NAD+/NADH redox pair as follows:

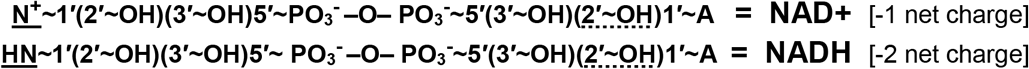

NAD’s adenine nucleotide holds one bit of information read by enzymes that recognize the absence (dot underlined hydroxyl group above) or presence of a phosphate group at the 2’ ribose carbon (dot underlined phosphate group below). This phosphorylation site is far removed from the nicotinamide base that serves as the electron-carrying moiety and is thus functionally independent of NAD’s redox state. While NADH is mainly used as a reducing agent in catabolic reactions and in the electron transport chain of oxidative phosphorylation, NADPH is mainly used as a reducing agent in anabolic reactions. We can diagram the oxidized and reduced forms of phosphorylated NAD as follows:

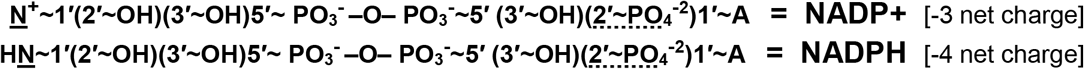

The ability to phosphorylate NAD allows cells to regulate NAD redox balances in two separate pools (+/-PO_3^-^_) for each membrane bound compartment (*e.g*., cytosol, mitochondrial matrix, stromal matrix of plastids, and peroxisomes). This enzymatic phosphorylation of NAD+/NADH is carried out exclusively by the NAD kinase (NADK) family. Members of the NADK family can differ by: **(*i*)** substrate specificity for NAD+ and/or NADH; **(*ii*)** donor source of the phosphate group, using either ATP and/or inorganic polyphosphate, poly(P); and **(*iii*)** subcellular localization. NAD(P)H is inferred to have been present in the last universal common ancestor (LUCA) of Bacteria, Archaea, and Eukarya (1–4, 8).

Despite the central role of NADK in the metabolism of cells since LUCA, and despite previous biochemical isolation of this enzymatic activity (9), the genes encoding it in model systems such as *E. coli* (*nadK*) and humans (cytosolic *NADK* and mitochondrial *NADK2*) were only identified recently in the post-genomic era (4, 6, 7, 10). How the human *NADK* and *NADK2* genes are related to other eukaryotic paralogy groups such as the three plant NADK genes (2, 3, 11), and how they are related to the *nadK* genes of various prokaryotic clades has not been determined.

Here, we curate and analyze *NADK* genes from several different eukaryotic and prokaryotic clades to identify the origin of eukaryotic *NADK* genes and their manner of evolution. We find that eukaryotic *NADK* gene evolution is episodic having been sensitive to major evolutionary transitions since the duplication of an *NADK* gene in early eukaryotic evolution. This ancient duplication established two clades of enzymes that ancestrally corresponded to cytosolic (“cyto”-clade) and mitochondrial (“mito”-clade) forms that are maintained in plants, most animals, and some protist groups. Other eukaryotic clades, such as chlorophytes or the clade represented by Holomycota + Amoebozoa have lost their mito-clade gene while duplicating and specializing a second cyto-clade gene for mitochondrial function (“c2m”). In contrast, most other protists have *NADK* genes that cluster in their own lineage-specific cyto-clades. Last, a few protist clades such as Choanoflagellata and Euglenozoa have more recently lost their mito-clade *NADK* gene while gaining an *nadK* gene from α-proteobacteria. Altogether these analyses show that there is a robust phylogenetic signal in the *NADK* sequences and in the particular repertoires of *NADK* genes.

## RESULTS

### A pair of duplicated *NADK* genes originated early in eukaryotic evolution

To identify the evolutionary origin of the eukaryotic NADK paralogs, we phylogenetically analyzed diverse eukaryotic groups together with the Archaea, the α-proteobacteria, and the cyanobacteria, which are the proposed origins of the ancestral eukaryotic host cell, its mitochondria, and of plastids, respectively, and additional prokaryotic groups (see Materials & Methods). Our explorations of the NADK tree space identify two ancient eukaryotic *NADK* sister clades of ancient unknown provenance (Fig. 1). We refer to these sister clades as the eukaryotic “cyto” and “mito” clades for the inferred ancestral function in an early or stem eukaryote clarity, and for reasons of clarity. This suggested convention bypasses the unrelated NADK numbering systems of vertebrates (*NADK1* and *NADK2*) and plants (*NADK1, NADK2, NADK3*). For example, human NADK2, which localizes to the mitochondria (10), is not orthologous to *Arabidopsis* NADK2, which localizes to the stromal matrix of chloroplasts (1, 2); human *NADK2* belongs to the mito-clade while *Arabidopsis NADK2* belongs to one of two plant cyto-subclades. While the main cytosolic enzymes of humans and plants (both named NADK1) belong to the “cyto” clade, the human mitochondrial enzyme NADK2 and the plant peroxisomal enzyme NADK3 belong to the “mito” clade (3, 10) (Fig. 1). This can be understood through comparative genomic data that suggests many nuclear-encoded mitochondrial proteins have been secondarily re-targeted to the peroxisome in some eukaryotes (12).

**Figure 1.**
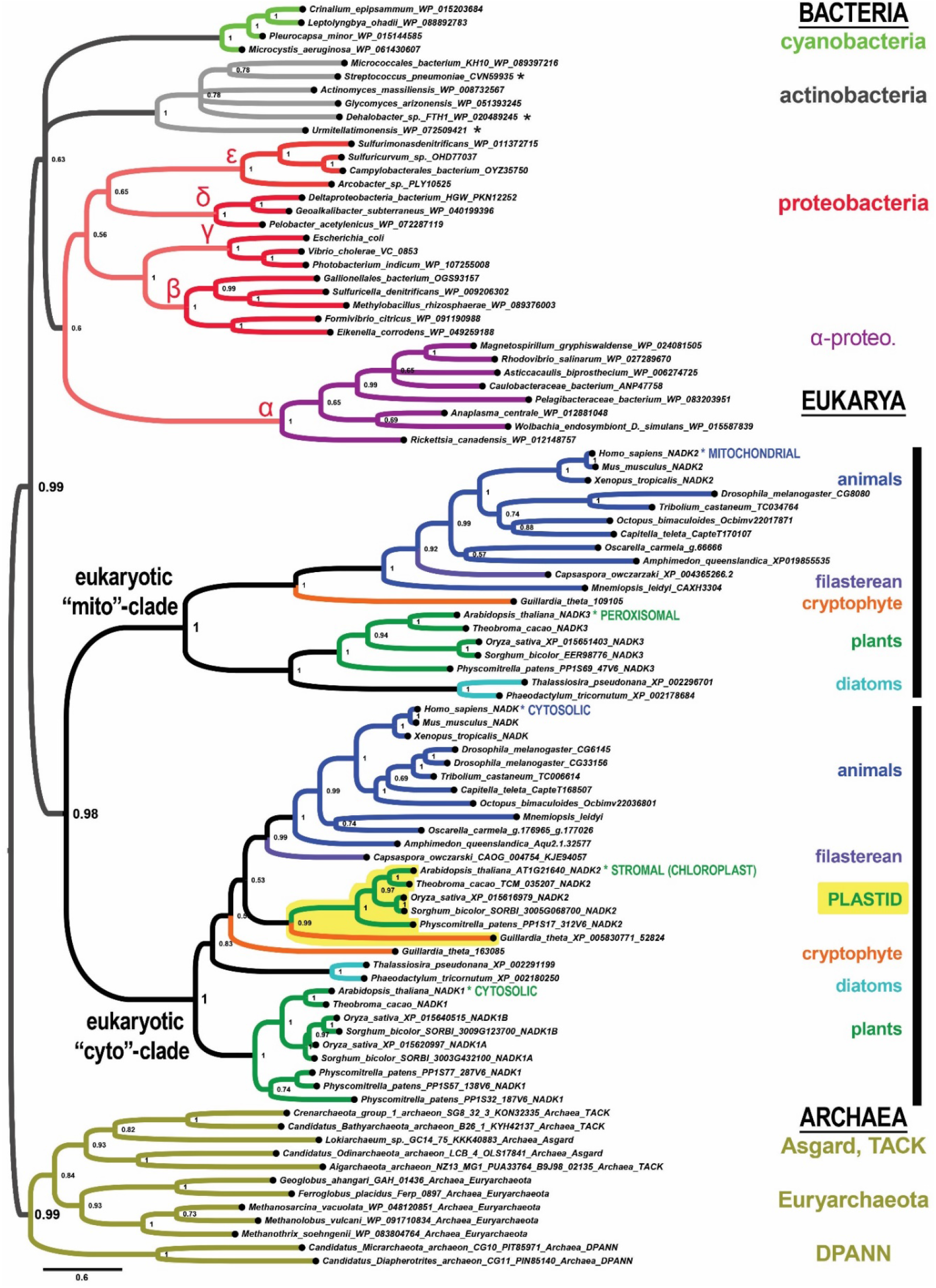
The *NADK* genes of extant eukaryotes originate from two ancient eukaryotic sister clades of uncertain provenance. Shown is a phylogenetic tree computed by Bayesian inference of *NADK* genes from the Bacteria (top clades), Eukarya (middle two clades labeled “mito” and cyto”), and the Archaea (bottom gold clade). Most animals (blue clade), plants (green clade), and a few other protist groups (labeled) have genes from both the mito and cyto subclades. Plants have genes from two cyto-subclades, one of which corresponds to the plant plastid-specific *NADK2* gene. The asterisked labels “MITOCHONDRIAL”, “CYTOSOLIC”, “STROMAL (CHLOROPLAST)” and others indicate lineages with specific experimental data on protein subcellular localization (see text). Node supports in this tree and all other trees represent posterior probabilities. Asterisks indicate sequences from species classified as Firmicutes but which we suspect are lateral gene transfers or misclassifications (also see Fig. S1).

A few major features characterize the ancient eukaryotic configuration of *NADK* genes carried by most plants and animals, and some eukaryotic protists (Fig. 1). The first feature is that the ancient eukaryotic nuclear genome encoded two eukaryotic paralogs that are sister clades to each other rather than to any one specific extant prokaryotic clade.

The second major feature is that *NADK* gene duplications predominantly occur in the cyto-clade with only one major exception specific to nematodes, which we describe in a separate section. Some of these duplications are associated with clade-defining evolutionary events (Table 1). For example, land plants and chlorophytes encode a plastid stromal matrix-specific paralog belonging to the cytosolic clade (yellow-highlighted clade in Fig. 1). Thus, the chloroplast *NADK2* genes of plants is derived from a duplication of the ancient cyto-clade eukaryotic gene rather than an endosymbiotic transfer from the cyanobacterial progenitor of the plastid.

**Table 1.**
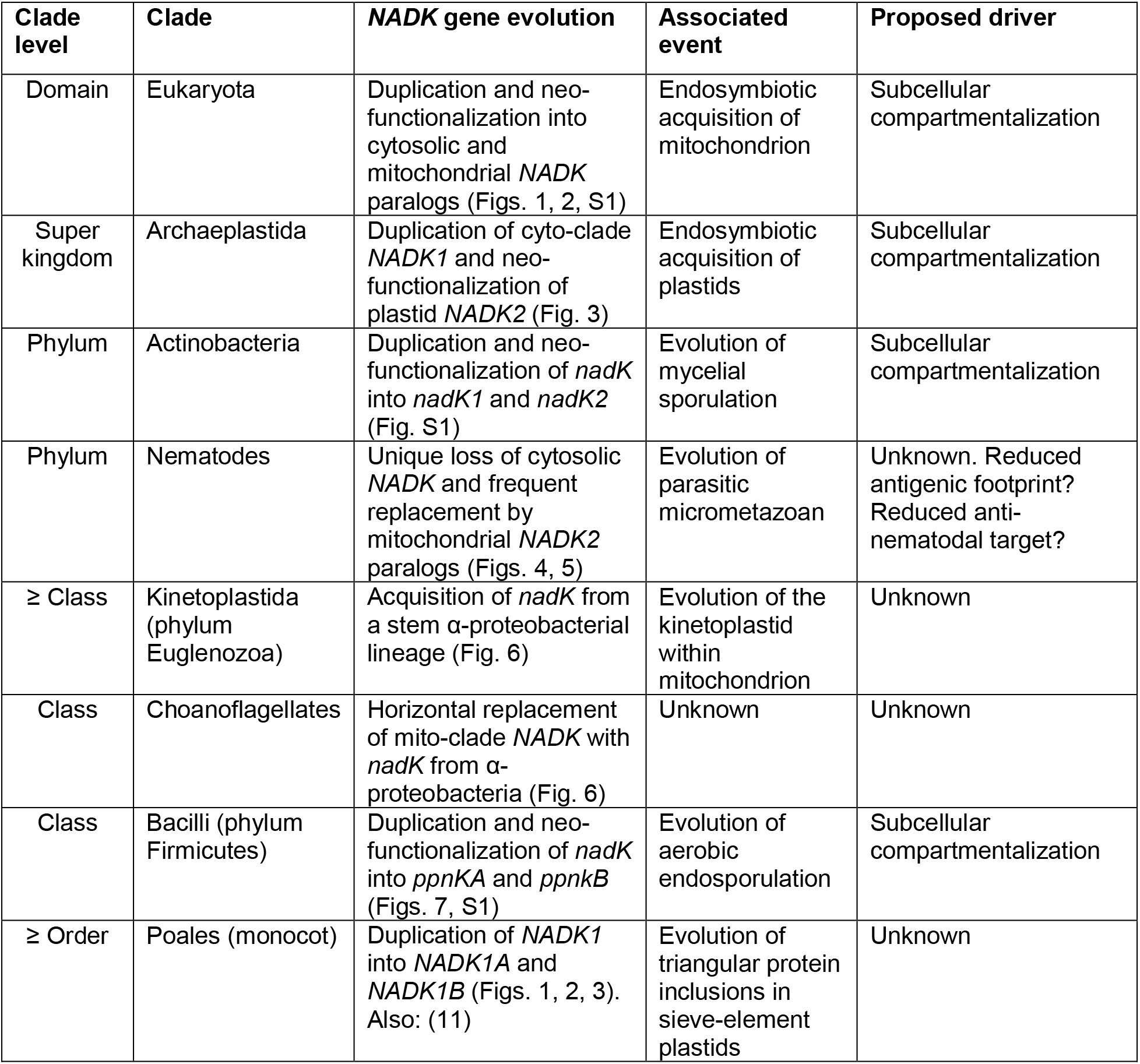
Episodic *NADK* duplications associated with clade-defining evolutionary transitions. Rows are ordered from highest to lowest clade levels. Gene duplications identified in a single species are not shown here.

The third major feature is that there is no consistent prokaryotic sister clade for the eukaryotic super clade. In our main Figure 1 tree (and other trees), one can see a highly supported Archaea composed of representatives from the Asgard, TACK, Euryarchaeota, and DPANN clades (Fig. 1 mustard-highlighted clade). A weakly supported Bacteria features highly-supported clades of representative cyanobacteria (light green), actinobacteria + firmicutes (light gray), and proteobacteria sub-clades (labeled with their Greek letter designations in the red highlighted clades in Fig. 1 and remaining figures). Furthermore, the α-proteobacterial *nadK* clade (the so-called purple bacteria), including both free-living (e.g., Rhodospirillales, Caulobacterales, and Pelagibacerales) and endosymbionts (Rickettsiales) is notably long-branched. Their NADK sequences share multiple derived α-proteobacteria-specific changes.

Some important eukaryotic clades follow a pattern wherein the ancestral mitochondrial NADK paralog is absent and apparently substituted by a neo-functionalized paralogous member of the cyto-clade. The fungal kingdom together with the unranked amoebozoans, and the chlorophyte division of plants (Viridiplantae) are such clades that appear to have lost their mito-clade *NADK* gene in connection with independent duplications of their cyto-clade *NADK* gene (Fig. 2, clades highlighted in yellow shading). For example, the chlorophytes *Chlamydomonas* and *Volvox* each have a plant-type *NADK1* gene (cytosolic form) and a plant-like *NADK2* gene (plastid form). However, unlike the embryophytes, including mosses (*Physcomitrella*) and angiosperms (dicots and monocots), and non-embryophyte streptophytes such as the charophyte *Klebsormidium nitens*, chlorophytes lack a detectable mito-clade *NADK* gene (plant *NADK3*). Instead, the chlorophytes have a third “cyto” clade (Fig. 2, bright yellow shaded clade).

**Figure 2.**
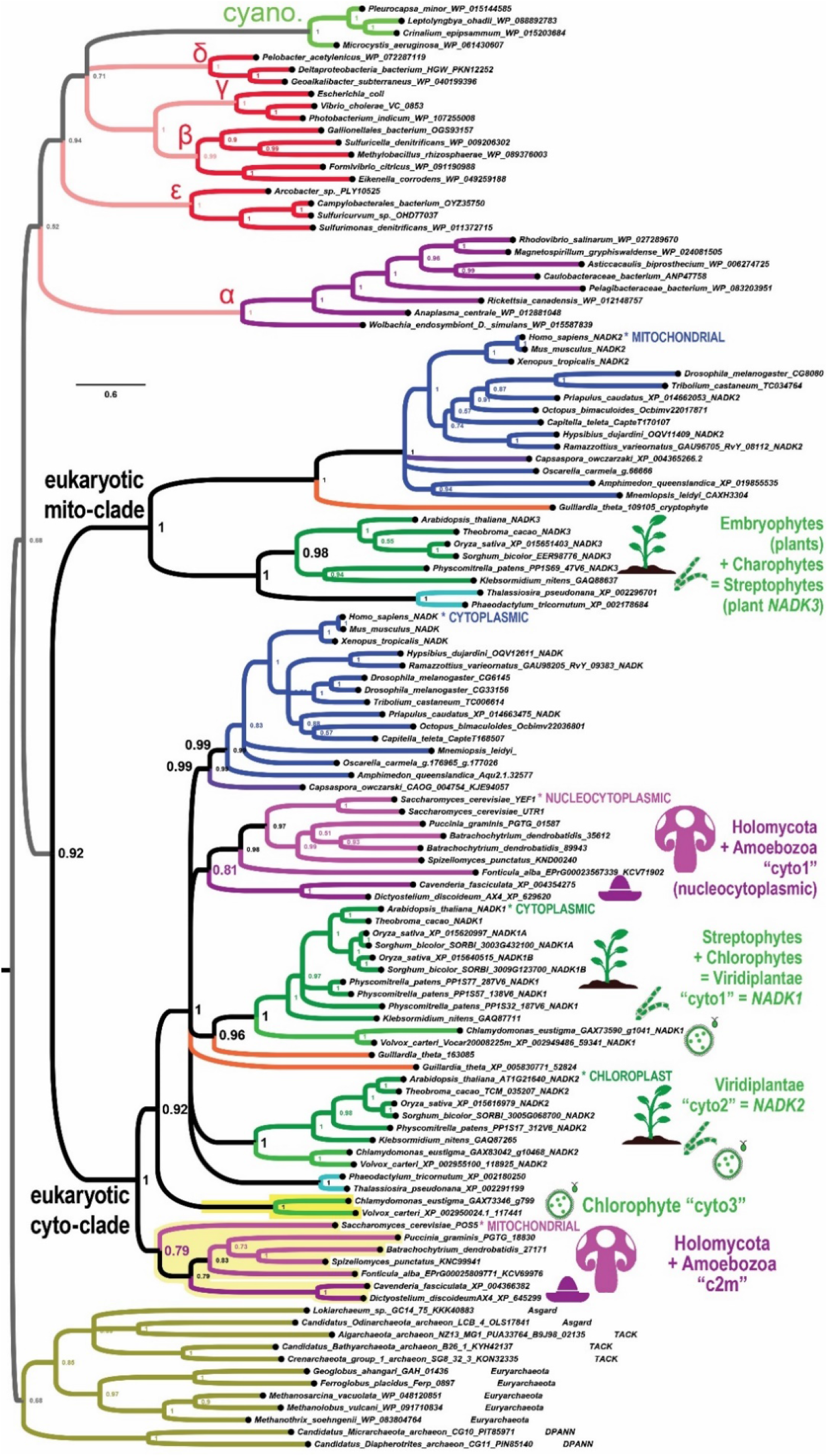
Fungi and Chlorophytes have lost mito-clade NADK genes while independently duplicating a cyto-clade gene. Plants (embryophytes), charophytes, and chlorophytes are each depicted with green icons, while Holomycota (fungi + *Fonticula alba*) and amoebozoans are each depicted with purplish pink icons. A pair each of the fungal groups of ascomycetes, basidiomycetes, and chytridiomycetes were chosen to represent the fungi. The extra cyto-clades for Holomycota + Amoebozoa and for chlorophytes are highlighted in yellow and are found at the bottom of the eukaryotic cyto-subclade.

Similar to the chlorophytes, the fungi and amoebozoans possess a clade of *NADK* genes grouping well within the cyto-clade and a more a basally branching duplicate cyto-clade that is not part of the mito-clade (Fig. 2 light yellow shaded clade). In *Saccharomyces cerevisiae*, duplicate genes (*YEF1* and *UTR1*) in the embedded cyto clade correspond to nucleocytoplasmic versions as expected, while the basally branching *POS5* gene corresponds to a known mitochondrial version (13) (“c2m”). Furthermore, the unusual, non-amoebozoan, cellular slime mold, *Fonticula alba*, which together with Fungi represent the Holomycota, appropriately follows the fungal pattern in its *NADK* gene repertoire.

The second basal “c2m” subclade of Holomycota + Amoebozoa is a sister-clade to the entire cyto-clade as opposed to being a sister-clade to the main cyto-clade containing *YEF1* and *UTR1* (Fig. 2). An interpretation of the paralogous fungal “c2m” clade requires that Holomycota and Amoebozoa are a monophyletic clade that excludes Holozoa. Or, if instead Holomycota and Holozoa form a monophyletic clade that excludes Amoebozoa, then one has to suppose the more complicated scenario that Holozoa has lost this additional “c2m” gene. Alternatively, these extra cyto-clades might represent divergent clades that have “fallen out” of the eukaryotic mito-clade.

We find that other eukaryotic protists harbor only cyto-clade *NADK* genes and frequently these have duplicated in their specific lineages (light purple highlighted clades in Fig. 3). Some of these are interesting because of the divergent evolutionary pathways they have taken with regard to their mitochondrion-related organelles (14). We summarize the results for each separate protist clade.

**Figure 3.**
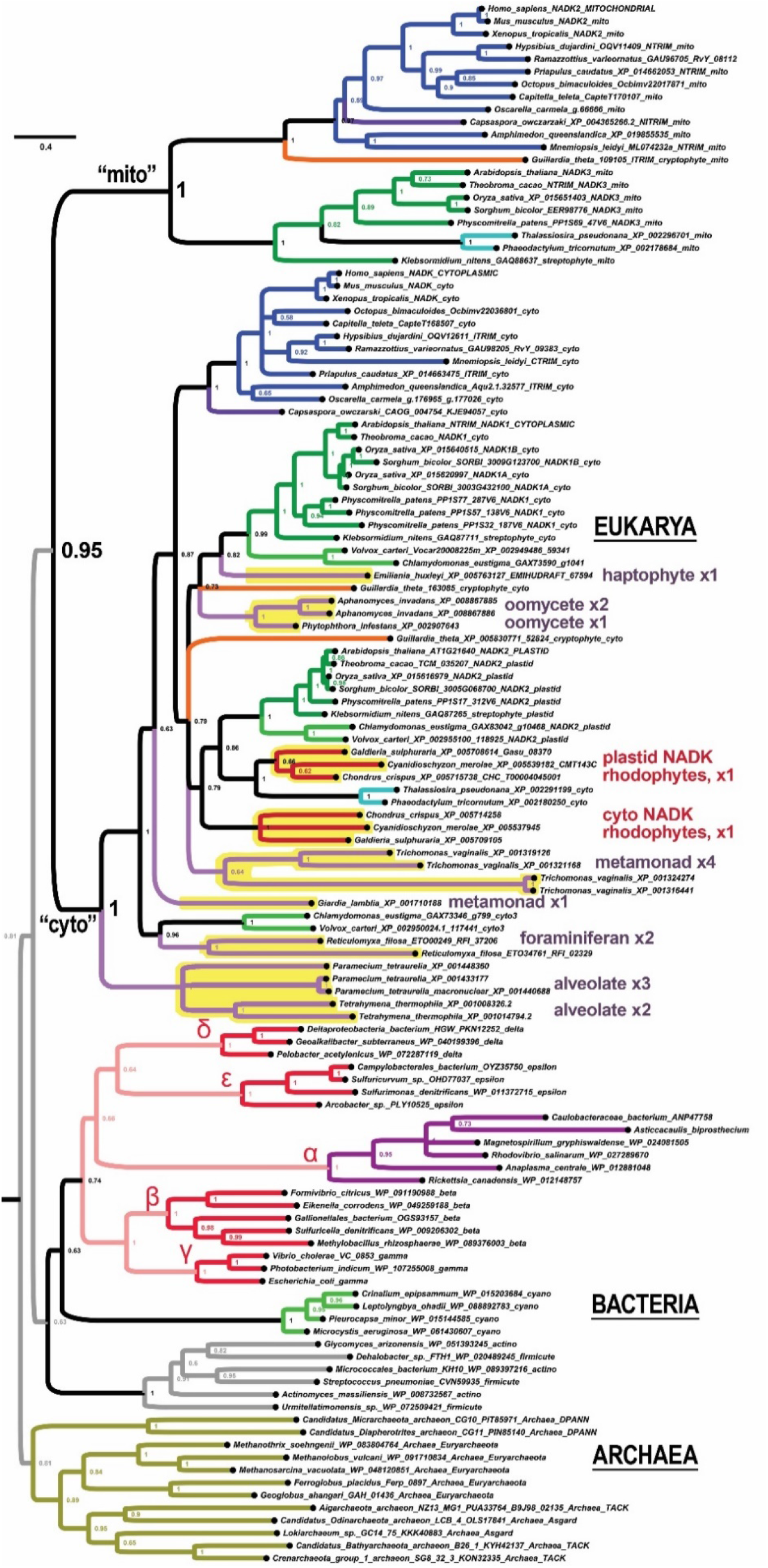
Many eukaryotic protists do not possess the mito-clade *NADK* gene of plants and animals. Unlike fungi, amoebozoans, and chlorophytes, all of which have additional *NADK* cyto-clade genes while missing a mito-clade *NADK* gene, many eukaryotic protist lineages that are also missing mito-clade *NADK* genes possess *NADK* cyto-clade genes that are only recently duplicated, if at all.

#### Alveolates

We find lineage-specific duplications for the alveolate *Paramecium tetraurelia*, and these form a sister clade to the lineage-specific duplications of the alveolate *Tetrahymena thermophile* (Fig. 3). The alveolate NADK sub-clade is sister to the rest of the eukaryotic cyto clade, all of which is sister to the eukaryotic mito clade.

#### Rhizaria

We find lineage-specific cyto-clade duplications for organisms within two different clades of Rhizaria: the phylum Cercozoa and the subphylum Foraminifera of phylum Retaria. In most trees, the cercozoan *Plasmodiophora brassicae* has a pair of lineage-specific duplications while the foraminiferan *Reticulomyxa filosa* also has a pair of lineage-specific duplications. These two rhizarian typically group together with the third set of cyto-clade genes of chlorophytes, although this is weakly supported (posterior probabilities between 0.51 and 0.83 in different trees).

#### Oomycetes

We find that the oomycete *Aphanomyces invadens* has a lineage-specific pair of duplicated cyto-clade genes that are most closely related to each other. This gene is sister to the single gene found for the oomycete *Phytophthora infestans*.

#### Metamonads

We find that the flagellated trichomonad parasite *Trichomonas vaginalis* has two pairs of duplicated cyto-clade genes for a total of four *NADK* genes. All four group together as three different duplications of different ages (cyto → cyto-A + cyto-B → → cyto-A1 + cyto-A2 and cyto-B1 + cyto-B2; see Fig. 3). These gene duplications may be consistent with the finding of one or more large-scale genome duplications in its recent past (15). In contrast, we find only one *NADK* gene for the flagellated diplomonad parasite *Giardia lamblia* but this is in the same sub-clade (typically) with the *Trichomonas* duplications.

The mitosome of the anaerobic *Giardia* is a derivative mitochondrial relic that no longer has a mitochondrial genome and no longer supports aerobic respiration (14, 16–20). The only known function of its mitosome is to continue supporting iron-sulfur cluster formation (16, 18, 20). Thus, the finding of a single cyto-clade *NADK* gene is consistent with the (inferred) loss of its mito-clade *NADK* gene.

*Trichomonas vaginalis* also features a derivative and non-conventional mitochondrial organelle (14). This double-cell walled organelle has been metabolically reprogrammed as a hydrogenosome that produces molecular hydrogen and ATP from the byproducts of glycolysis in the cytosol. The hydrogenosome continues to support several metabolic synthesis pathways that would require NADPH (21). Thus, it would make sense that *Trichomonas* continues to harbor multiple genes in contrast to the single gene of *Giardia*.

#### Rhodophytes

Rhodophyta and Viridiplantae together comprise the Archaeplastida, which are supposed to have descended from the same primary endosymbiotic event between a eukaryotic protist and a cyanobacteria. We find that three species of rhodophytes from three genera each have a pair of cyto-clade *NADK* genes grouping into two separate rhodophyte sub-clades (Fig. 3). One of these rhodophyte sub-clades is sister to the Stramenopiles cyto-clade and together these are sister to the *NADK2* plastid clade of Viridiplantae. This grouping is thus consistent with Archaeplastida. The second rhodophyte cyto-clade is also sister to the Archaeplastida *NADK2* clade, which would be partially consistent with an ancient cyto duplication into the plant *NADK1* and *NADK2* gene clades.

#### Haptophytes

We find only a single identifiable *NADK* gene for the haptophyte *Emiliania huxleyi* (Fig. 3). This gene groups basally with the plant NADK1 clade consistent with acquisition of photosynthesis from a higher-order endosymbiotic event.

### Unique fractal evolution of mito-clade *NADK* in nematodes

We find that the metazoan phylum of nematodes presents a unique pattern *NADK* evolution compared to all other eukaryotes. Whereas we observed all eukaryotes to encode either zero or one mitochondrial *NADK* gene (the *“NADK2”* gene of animals, which is orthologous to the “*NADK3*” gene of plants), the ancestral nematode simultaneously lost its cyto-clade *NADK* gene, while duplicating its mito-clade gene (Fig. 4). We can infer that this event occurred specifically within the nematode clade because we are able to find both cyto- and mito-type *NADK* genes in insects (the dipteran *Drosophila* and the coleopteran *Tribolium*) as well as in tardigrade genera (*Hypsibius* and *Ramazzotius*) and in the priapulid worm *Priapulus caudatus* (Fig. 4). In contrast, we only find nematode *NADK* genes within the mito-clade (Fig. 4 and Fig. 5). All of the ecdysozoan mito-clade *NADK* genes, including those of nematodes, form a well-supported sister group to lophotrochozoan genes in a well-supported protostome clade (Fig. 4).

**Figure 4.**
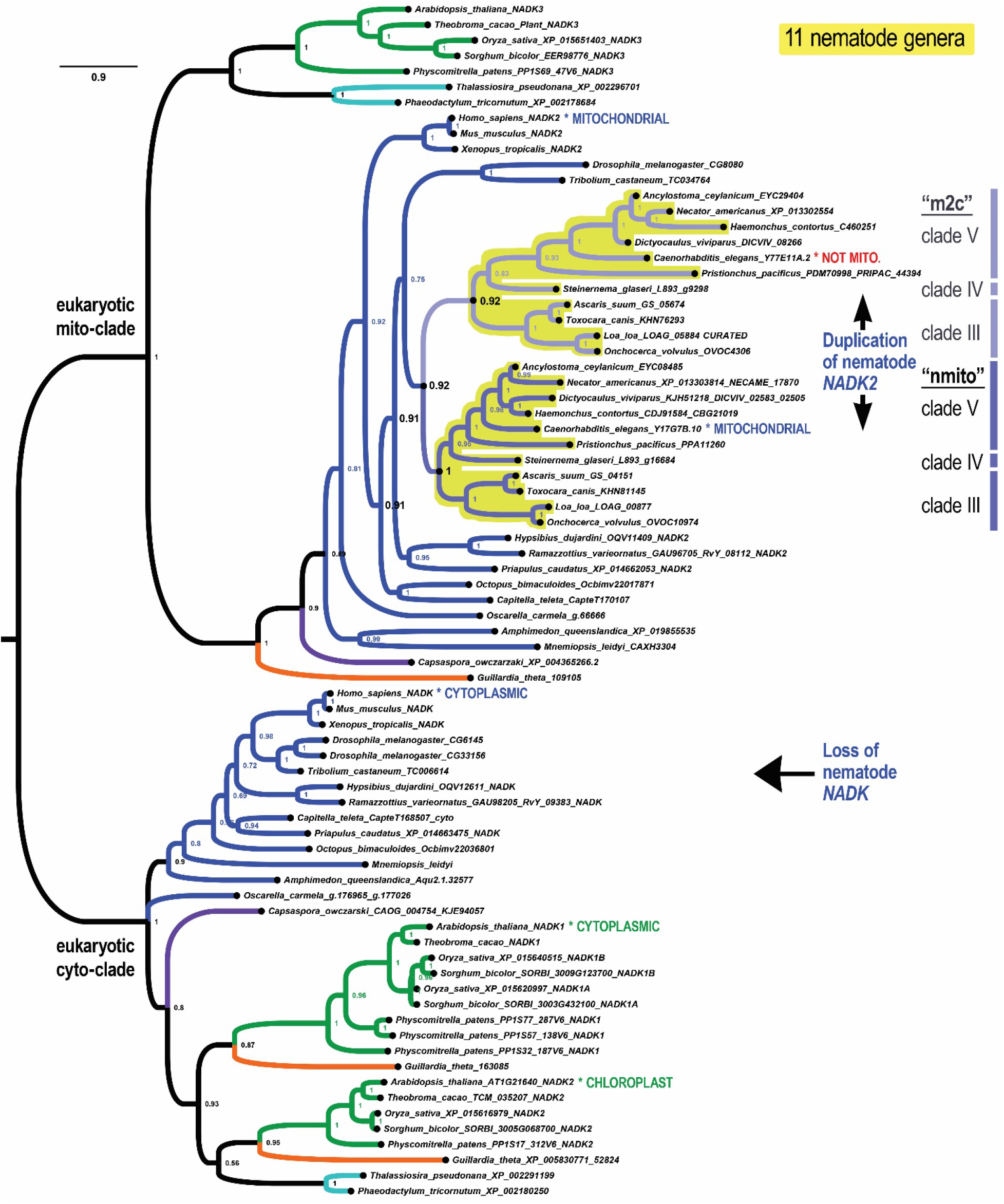
An ancient duplication of *NADK* characterizes Nematoda. All nematodes have missing cyto-clade genes, which was lost in connection with a duplication of the ancestral nematode mito-clade gene. Shown is a phylogenetic tree of *NADK* genes from 11 different nematode genera (highlighted in yellow) that have the ancient nematode duplication of its mito-clade gene into an “nmito” paralog, which encodes an enzyme localized to the mitochondrion in *C. elegans*, and an “m2c” paralog, which no longer localizes to the mitochondria in *C. elegans*.

**Figure 5.**
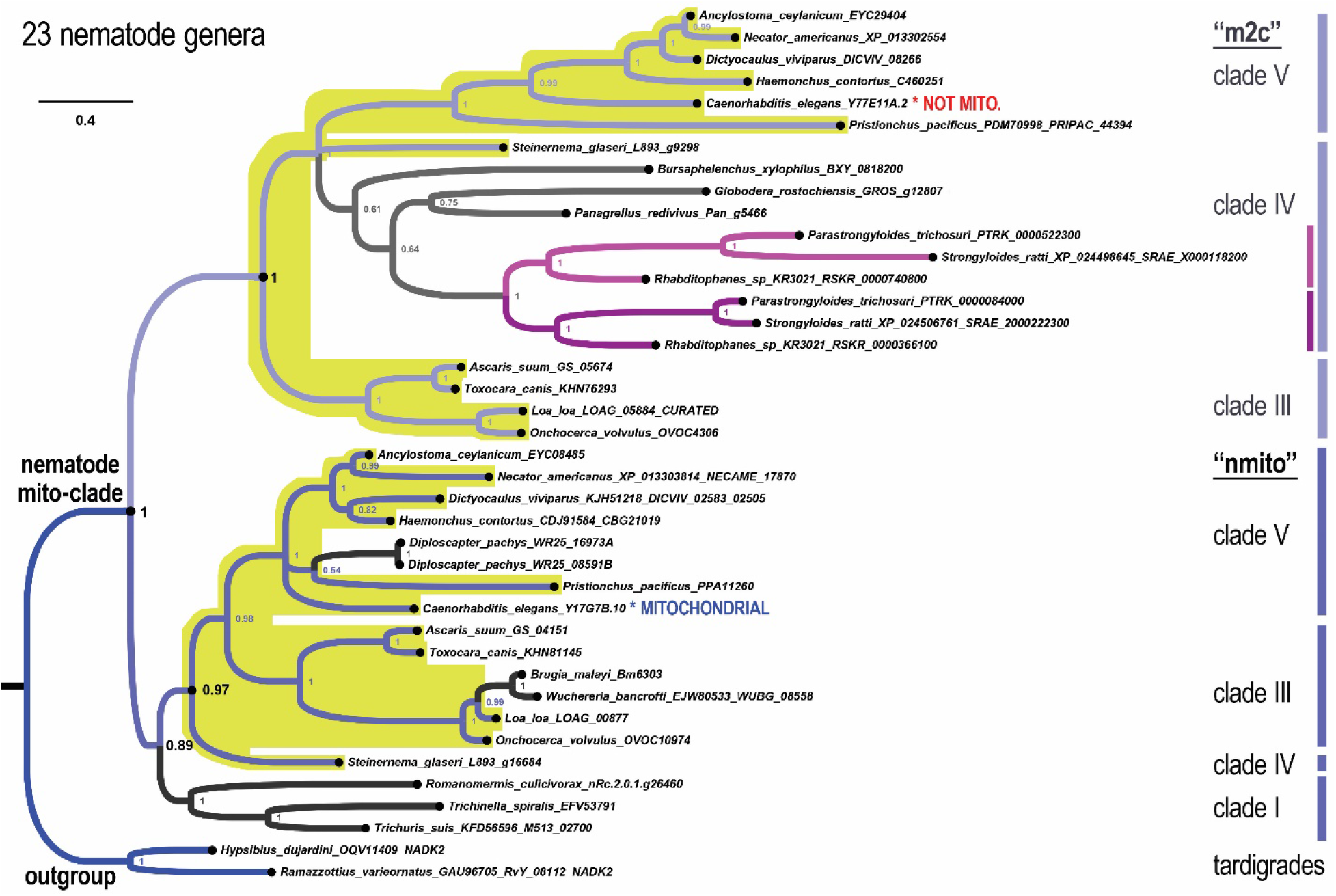
A fractal-like evolutionary pattern characterizes *NADK* evolution in the nematode phylum. Shown is a phylogenetic tree of the nematode mito clade with additional nematode genera that display more recent duplications such as the clade IVb (purplish lineages) in the m2c-subclade and *Diploscapter* genes in the nmito-subclade of clade V nematodes. Other lineages, such as clade I nematodes appear to have only one identifiable gene within one of the two nematode subclades.

Because the slower-evolving *NADK2* paralog of *C. elegans* encoded by Y17G7B.10 is known to be mitochondrially-localized while the faster-evolving Y77E11A.2 is not (22), we refer to the genes in the nematode-specific clade containing the former as the “nmito”-subclade (for “nematode mitochondrial”) and the genes in the nematode-specific clade containing the latter as the “m2c”-subclade (for “mito-to-cyto”). The nmito and m2c clades are highly-supported sister-clades in the highly-supported nematode-specific mito-clade (Fig. 4 and Fig. 5).

Current understanding of nematode phylogeny divides the phylum into a series of numbered clades corresponding to the Dorylaimia (clade I), the Enoplia (clade II), and the Chromadoria (composed of clades III, IV, and V) (23, 24). Figure 4 depicts *NADK* genes from nematodes in clades III, IV, and V that have retained both the nematode nmito-type and m2c-type genes. In contrast, Figure 5 depicts additional *NADK* genes from nematodes that we observe to have diverged from the ancestral nematode pattern. We now summarize an interesting fractal-like pattern of *NADK* evolution across the various nematode clades for which we had available data (all except clade II/Enoplia). We then speculate on the significance of this evolutionary pattern in the Discussion section.

#### Clade V (Rhabditina)

The overwhelming majority of clade V genera that we analyzed (6/7), and which includes the well-studied *Caenorhabditis elegans* species, have both an nmito- and an m2c-type *NADK* gene (Fig. 4). The single exception is for the pair of genes from *Diploscapter*, whose genes represent a lineage-specific duplication of an nmito-type gene along with an inferred loss of its m2c gene (Fig. 5).

#### Clade III (Spirurina)

Like clade V, the majority of clade III genera that we analyzed (4/6) have both an nmito- and an m2c-type *NADK* gene (Fig. 4). The exception is for the genes from *Brugia malayi* and *Wucheria bancrofti* for which we were only able to find a single nmito-type gene from each. For example, we were not able to find credible matches to m2c-type genes even by using m2c protein sequences to search these genomes using TBLASTN (protein to translated nucleotides) and relaxed parameters (PAM250 and BLOSUM45 substitution matrices with relaxed gap penalties).

#### Clade IV (Tylenchina)

Except for the nmito and m2c genes from *Steinernema glaseri* (Fig. 4), the rest of the clade IV sequences, which pertain to 6 additional genera, belong exclusively to the m2c-type clade. Interestingly, three of these genera have a shared ancestral duplication of their m2c gene. Thus, on the basis of this m2c duplication alone, we can phylogenetically place the *Parastrongyloides, Strongyloides*, and *Rhabditophanes* genera in a single “IVb” sub-clade. All of the remaining clade IV genera (4/7 genera) branch basally and are not resolved as a single monophyletic clade using just these *NADK* sequences.

#### Clade I (Dorylaimia)

Last, we were only able to find *NADK* genes from the nmito-clade for three genera of clade I nematodes, all of which are highly derived parasitic nematodes of mammals (*Trichinella* and *Trichuris*) or insects (*Romanomermis*) and one of which is an intracellular parasite (*Trichinella*) (Fig. 5). These were grouped together with a posterior probability of 1 and as a sister-clade to the remaining nmito genes with a posterior probability of 0.89 (Fig. 5). Thus, this topology does not support that these sequences are a sister-clade to a super-clade of nmito and m2c sub-clades, but rather that clade I nematodes lost their m2c genes. Nonetheless, we cannot completely rule out that the ancestral nematode duplication that produced the nmito and m2c paralogs occurred during early nematode divergence.

### Loss of mito-clade *NADK* with horizontally-transferred α-proteobacterial *nadK*

We find three eukaryotic clades/lineages have lost mito-clade genes while simultaneously gaining genes from different lineages of α-proteobacteria. The first example concerns the choanoflagellates *Monosiga brevicollis* and *Salpingoeca rosetta*, which encode cyto-clade NADK sequences that are most closely related to each other in a holozoan clade that includes metazoans and the filozoan Capsaspora (Fig. 6 “choano-cyto”). However, the choanoflagellates lack NADK sequences in the mito subclade (Fig. 6 “choano-mito loss”). Instead of having canonical holozoan mito-clade genes, choanoflagellates harbor *nadK* genes horizontally acquired from the α-proteobacteria (Fig. 6 “choano-alpha”). That this is a horizontal gene transfer (HGT) is further supported by the presence of amino acid sequences only shared by the α-proteobacteria NadKs. HGT is also implied given the presence and position of the more ancient NADK paralogy groups shown in Figure 1.

**Figure 6.**
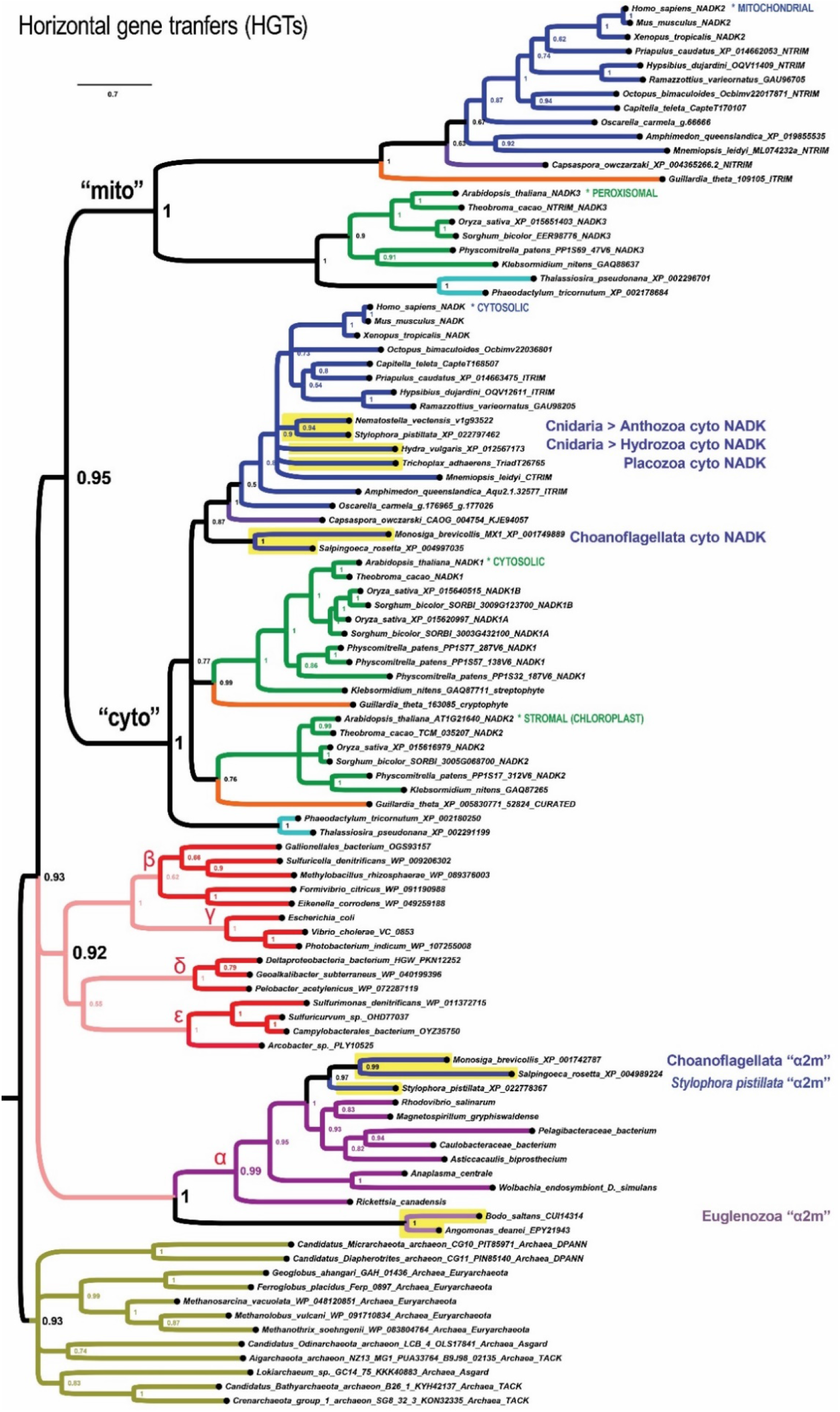
Rarely, eukaryotic lineages replaced mito *NADK* genes with horizontal transfer of α-proteobacterial *nadK*. Choanoflagellates, euglenozoans, and the cnidarian *Stylophora pistillata* eukaryotic mito-clade *NADK* genes while also possessing nadk genes apparently acquired from different lineages of α-proteobacteria. These apparent cases of HGT thus appear to be “α2m” replacements of mito-type NADK.

We find another apparent case of mito-clade replacement with α-proteobacterial HGT with one of the two anthozoan species of cnidarians, *Stylophora pistillata* (Fig. 6). All other cnidarians that we analyzed, including the anthozoan *Nematostella vectensis* and the hydrozoan *Hydra vulgaris* have only a single gene each within the metazoan cyto clade. A similar case of a single cyto *NADK* gene characterizes the placozoan *Trichoplax adhaerens* (Fig. 6).

Last, we find that the free-living, euglenozoan excavate *Bodo saltans* and another euglenozoan excavate that is an obligate parasite of some insects, *Angomonas deanei*, have *nadK* genes that form a sister clade with the α-proteobacteria. However, unlike the choanoflagellates and the cnidarian *Stylophora pistillata*, the euglenozoans possess only this single *NADK* gene (Fig. 6). Nonetheless, the euglenozoan genes are consistently paired with each other across different trees as are the choanoflagellate choan-alpha genes consistent with these two eukaryotic clades being associated with independent horizontal gene transfers (HGTs) of proteobacterial and α-proteobacterial *nadK*, respectively.

### Independent duplications of *nadK* in sporulating bacteria

We have shown that eukaryotes originated with two ancient *NADK* paralogs of uncertain provenance in relation to prokaryotes. While it has been assumed that prokaryotes typically only have a single *nadK* gene, we desired to test this assumption systematically for comparison to eukaryotes. To investigate whether bacterial and archaeal cells ever have more than one *nadK* gene, we first searched the ComparaEnsembl orthology calls (25). For example, we identified the number of genes per species on bacteria.ensembl that are orthologous to the single *nadK* gene in *Escherichia coli*. Of the 77 bacterial and 23 archaeal species with *nadK* genes analyzed by EnsemblCompara, only 3 bacterial species have multiple *nadK* genes identified. Two of these species were from the Firmicutes phylum, *Bacillus subtilis* and *Listeria monocytogenes*, while the third was from the actinobacterium *Streptomyces coelicolor*. Each has a pair of *nadK* genes.

To determine the relationships between the gene duplicates of *Bacillus, Listeria*, and *Streptomyces*, we computed phylogenetic trees of bacteria using additional species of Firmicutes and Actinobacteria taking care to sample genes coming from the same strains (Fig. 7 and Fig. S1). We find that these duplicated *nadK* paralogs stem from two independent duplications in the bacterial phyla of Firmicutes and Actinobacteria (Fig. 7 and Fig. S1). For the duplication within Firmicutes, we find that we can place this duplication as occurring within the stem lineage leading to the class Bacilli because the *nadK* genes in the sister class of Clostridia form a sister clade to the Bacilli paralogs (Fig. 7). Thus, many species within the class Bacilli have two genes for *polyphosphate/ATP-dependent NAD kinase A* (*ppnKA*) and *ppnKB* as named in *Bacillus subtilis* (Fig. 7). The duplication in *Streptomyces* is shared widely within Actinobacteria because the paralogs *“nadK1”* and *“nadK2”* (so named here to distinguish them from the independent duplication in Bacilli) can be found in multiple orders of Actinobacteria: Streptomycetales (*Streptomyces* and *Streptacidiphilus*), Pseudocardiales (*Amycolatopsis*), and Propionibacteriales (*Kribbela*) (Fig. S1).

**Figure 7.**
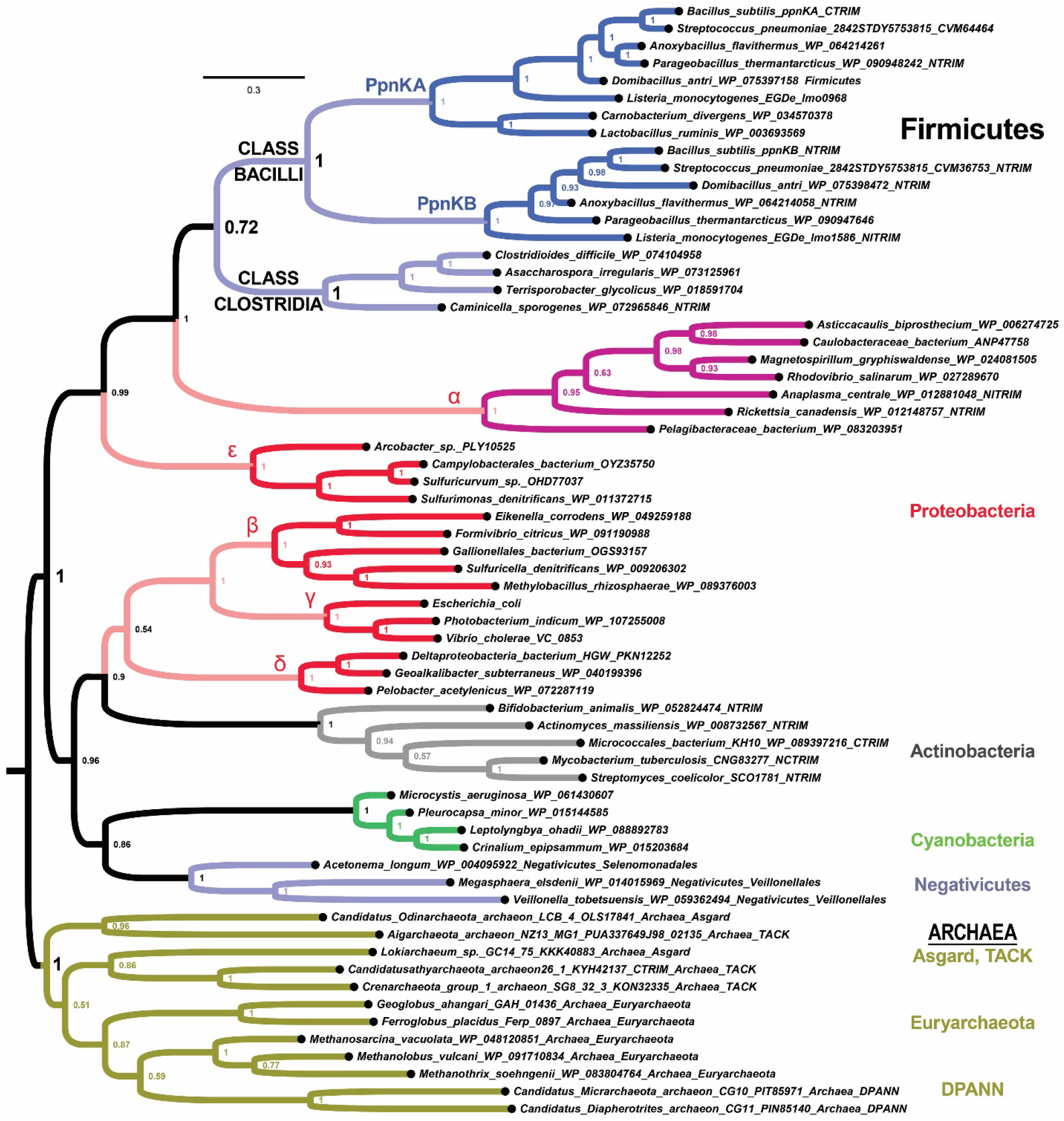
A bacterial duplication of *nadK* in the Firmicutes class of Bacilli. Identification of a duplication of *nadK* into *ppnKA* and *ppnKB* in the occurred after the Bacilli class split with its sister class of Clostridia. The Bacilli class of Firmicutes corresponds to aerobic endospore formers while the Clostridia class corresponds to anaerobic endospore formers. See Figure S1 for a more complete tree that includes the independent duplications in Actinobacteria and Eukaryotes.

Current taxonomic classification based on 16S rRNA splits the Firmicutes class Bacilli into the paraphyletic order Bacillales and the monophyletic order Lactobacillales (26). We find that some Lactobacillales genera (*Lactobacillus* and *Carnobacterium*) appear to have lost *ppnKB* while others (such as *Streptococcus*) have not (Fig. 7 and Fig. S1). Thus, this *bona fide* paralogous duplication of *nadK* into *ppnKA* and *ppnKB* occurred in the stem Bacilli lineage after the split from class Clostridia. An alignment of different Bacilli NadKA and NadKB proteins shows that there are several residues throughout the enzyme that are evolutionarily differentiated in the NadKA clade and/or the NadKB clade (Fig. S2). Altogether, our results show that the independent bacterial duplications in the Firmicutes class *Bacilli* (Fig. 7 and Fig. S1) and the Actinobacteria phylum (Fig. S1) is unrelated to the ancestral duplication in LECA.

To address whether anything is known about differential regulation of *ppnKA* and *ppnKB* in the well-studied *Bacillus subtilis*, we searched a transcriptomic profile generated over hundreds of conditions (27). Growth profiles in LB, LB + glucose, and M9 minimal media indicate that *ppnKA* has its highest expression levels during exponential growth when *ppnKB* was having its lowest expression levels, while *ppnkB* was having its highest expression levels during stationary phases when *ppnKA* was having its lowest expression levels (27). Confluent plate growth, induction of swarming, and sporulation (1 hour post initiation of sporulation) are also separate conditions that caused the highest observed expression levels for *ppnkB* but not *ppnKA* (27). We discuss the significance of these transcriptomic profiles below.

## Discussion

Our results show that *NADK* genes from all domains of life possess a potent phylogenetic signal in the genes themselves and in the series of duplications associated with major clade-defining events in cellular evolution. We identify at least five major duplications associated with the highest taxonomic levels (Table 1). The first major duplication is one associated with the acquisition of the mitochondrion by the latest eukaryotic common ancestor (LECA). We find that the ancient cytosolic and mitochondrial forms of *NADK* were already present as sister paralogs in LECA. Furthermore, we find that these two ancient eukaryotic paralogs could not be definitively assigned either to proteobacteria or to Archaea and are of unknown provenance. Thus, all we can say is that LECA originated with two ancient *NADK* genes specialized for subcellular compartmentalization. If we assume the α-proteobacterial origin hypothesis for mitochondria (28–30), then we must also conclude that the proto-eukaryotic host *NADK* gene duplicated, and then neo-functionalized and replaced the original mitochondrial *nadK* gene. Alternatively, if the eukaryotic mitochondrion is not descended from proteobacteria, then either the host or endosymbiont *nadK* gene could have duplicated in the stem-eukaryote.

A second major duplication event is associated with the higher order endosymbiotic acquisition of plastids in Archaeplastida (Table 1). This occurred by duplication of the cytosolic *NADK* gene of plants (embryophytes), charophytes, rhodophytes, diatoms, and likely cryptophytes. Thus, despite multiple examples of eukaryotic endosymbioses including the mitochondrial endosymbiosis of LECA, the primary and higher order endosymbioses of plants and cryptophytes, host gene duplication and neofunctionalization has predominated over endosymbiotic gene transfer.

We find other independent duplications at the phylum and sub-phylum (class) level in Bacteria and these are potentially interesting for their relationship to membrane compartmentalization associated with mycelial sporulation (Actinobacteria) or aerobic endosporulation (the Firmicutes class of Bacilli). We discuss these details further below.

Last, a striking duplication pattern is seen in the phylum of nematodes, which are unique among eukaryotes in having lost the main cytosolic NADK gene (Fig. 4 and Fig. 5). This is the converse situation to many eukaryotic protist clades (Fig. 2 and Fig. 3) in which the mitochondrial *NADK* gene is lost and apparently replaced with duplications of the cytosolic gene. We discuss this special finding in nematodes and its possible significance in relation to nematode biology next. We also propose gene names for the *Caenorhabditis* and *Drosophila* paralogs, all of which are currently unnamed, based on the paralogy relationships (Table S1).

We also found three rare groupings of eukaryotic *NADK* genes with α-proteobacteria: (i) choanoflagellates (*Monosiga brevicollis* and *Salpingoeca rosetta*), (ii) the anthozoan cnidarian *Stylophora pistillata*, and (iii) euglenozoans (*Bodo saltans* and *Angomonas deanei*) (Fig. 6). One possible explanation for the first two, which represent groupings of genes in addition to standard cyto-clade genes possessed by those species, or for all three, is that they represent the true affinity with the proposed clade of mitochondrial ancestry (28, 29). However, given that these sequences group within α-proteobacteria uniquely (*i.e*., mostly separately) and unlike those of the vast majority of eukaryotic *NADK* genes, we interpret these results as gene replacements by recent horizontal gene transfer from α-proteobacterial endoparasites. Frequent HGT from intracellular endosymbionts from this clade may have masked a more ancient provenance for other genes as well. This latter interpretation would be consistent with evidence discounting the α-proteobacterial hypothesis for mitochondria (31, 32).

### On the unprecedented loss of cytosolic *NADK* in nematodes

Nematode *NADK* evolution is fractal-like in that mito-clade gene duplications accompany cyto clade and mito subclade gene losses at multiple time scales. First, like the inferred ancient duplication of eukaryotes that produced the cyto-clade and mito-clade *NADK* genes, a nematode progenitor underwent an ancestral duplication of its mito-clade gene. This occurred in connection with a loss of its ancestral cyto-clade *NADK*, which is otherwise retained in diverse ecdysozoan phyla that we analyzed. Furthermore, there is evidence that the faster-evolving m2c paralog now serves the cytosolic role, while the slower-evolving nmito paralog retains the mitochondrial role. Of the 23 genera from four nematode sub-clades that we analyzed, 11 genera, or almost half, retain both of the ancestral nematode duplications (nmito and m2c types).

Second, four nematode genera have lost one of the two ancient nematode duplications, while undergoing a duplication of their remaining mito-clade gene. This occurred during clade IV divergence in the sub-clade IVb ancestor and more recently in the *Diploscapter* lineage within clade V. Thus, the majority of nematode genera analyzed (15/23) encode two mito-clade genes related to either an ancestral nematode-, a sub-clade IVb-, or a *Diploscapter-* specific duplication event. No other eukaryotic group has duplicated their mito-clade genes, much less repeatedly as in nematodes.

Whatever the reason for the nematode evolutionary pattern, the nematode case definitively establishes that loss of one eukaryotic *NADK* gene in an organism is frequently accompanied by duplication of another existing *NADK* gene in its repertoire. The nematode case for this apparent compensatory turnover of paralogs is definitive because the nematode nmito and m2c paralogs are sister clades that are deeply embedded in the eukaryotic mito-clade at the appropriate place within Metazoa and furthermore within Ecdysozoa. Thus, it is unlikely to be a long-branch attraction artifact of nematode cyto and mito clades. This interpretation is further supported by lineage-specific nematode duplications continuing to occur within these well-supported sister clades.

Nematodes are a micrometazoan phylum specialized for parasitism of multicellular plants, animals, and fungi. Greater than one-half of the named species have such lifestyles and ectoparasitic and endoparasitic forms have repeatedly evolved in different clades and across plant and animal Kingdoms (23). Endoparasitic forms also include both intra-tissue parasites as well as intracellular parasites (e.g., *Trichinella spiralis* larvae invade mammalian muscle cells). Last, even the non-parasitic forms are frequently found to be commensalists of other animals. Thus, we propose that one hypothesis for this persistent pattern of gene evolution in nematodes is that nematodes evolved stealth antigenic footprints and/or attenuated anti-nematodal targets for host defense pathways (33). Whatever the case, some specific evolutionary driver likely underlies the unique and recurrent pattern of *NADK* gene loss in nematodes.

### Evolution of membrane compartmentalization in sporulating bacteria

Given the *B. subtilis* expression profiles of *ppnkA* (exponential growth) versus *ppnkB* (stationary phase, swarming, confluence, and sporulation), the *nadK* duplication in Firmicutes is highly significant (Fig. 7 and Fig. S1). It is thought that the bacterial process of endospore formation arose only once in an ancestor of the Firmicutes, with the class Bacilli representing the aerobic endospore formers and the class Clostridia representing the anaerobic endospore formers (34).

Interestingly, the actinobacteria *Streptomyces* is thought to have evolved a type of sporulation (but not the endospore formation pathways of Bacilli) through independent mechanisms (34), and this also is coincident with a duplicated *nadK* gene in this clade (Fig. S1). In *Streptomyces coelicolor, nadK1* appears to be associated with cell growth (anabolic pathways) and sporulating stages in both liquid and (mycelated) solid cultures, while *nadK2* appears to be highest in sporulated spores following the mycelated sporulation in solid cultures (35, 36). Thus, this duplication in Actinobacteria might also be associated with NADK neofunctionalizations associated with mycelial-type sporulation pathways of Actinobacteria (37).

These results suggest that, like the process of membrane compartmentalization in Eukaryotes, the process of endospore formation or some aspects of sporulation relating to bacterial cell envelopes in general (38), may engender a need for different NadK enzymes. Altogether, these results suggest that eukaryotes may have evolved from a prokaryotic ancestor that evolved duplicate *nadK* genes in connection with membrane-bound compartmentalization associated either with endosymbiont recruitment and/or processes analogous to bacterial sporulation.

Last, one outstanding aspect of our results is the persistent affinity between the NadK sequences of Firmicutes and α-proteobacteria, which frequently are sister-clades (*e.g*., Fig. 7 and Fig. S1). One likely possibility is that this is the result of long-branch attraction. However, an alternative explanation not yet ruled out is that an α-proteobacterial ancestor acquired an NadK-encoding gene from the stem-Firmicutes lineage. In any case, our results indicate that the origin of the paralogous eukaryotic NADK cyto and mito clades are nonetheless of ancient and unknown provenance.

## Methods

### Protein alignment

We constructed amino acid alignments using the MUSCLE algorithm option the MEGA7 program (39, 40). Unique extensions in the N- and C-termini or unique internal loop insertions were trimmed. In the FASTA headers such sequences are labeled “NTRIM”, “CTRIM”, and “ITRIM” or combinations thereof (e.g., “NICTRIM”). We also extended partial automatic (hypothetical) gene predictions by consulting the genomic sequence. These sequences are typically labeled “CURATED” after the gene prediction ID in the FASTA header. For alignment, we mostly used the MEGA7 MUSCLE default parameters with adjustments in the gap existence and gap extension parameters. In preliminary explorations of tree space, while the NADK protein data set was being built up, the gap existence and gap extension parameters were left at default (−2.8 and 0.0, respectively). In later refinement of tree space, these were increasingly adjusted as follows. To adjust the ragged N-terminal and C-terminal prokaryotic subset of the data, the gap existence was adjusted down to −1.0. After aligning and trimming these sequences, these were then realigned with the full data set (i.e., with the eukaryotic clades included) using a gap existence of −2.0 down to −1.8 or − 1.5 (depending on the taxa) and a gap extension of −0.02. The specific values used will be listed for each representative tree depicted in the study.

### Phylogenetic analysis

All phylogenetic analyses were conducted using the 64 bit serial version of MrBayes on different Windows machines (41–44). Metropolis coupling (MC-MCMC) with two “nchains”, a temperature of 0.08, and double precision operations (BEAGLE library) were all enabled. In early analyses, we tested mixed protein substitution models with 25% burn-in generations in prokaryotic + eukaryotic trees, and in prokaryotic-only and eukaryotic-only trees, but as we kept getting the WAG substitution model in later analyses, we switched to using a fixed (WAG) substitution model and gamma-shaped frequency model with some invariant residues (“invgamma”)(45). Convergence of runs was evaluated by the average standard deviation of split frequencies, and the trees shown had values less than 0.01 (unless otherwise stated) after 300k to 2 M generations (typical run was 1.2 M generations long). Trees were rendered and decorated using the FigTree 1.4.3 tree figure drawing tool.

## Acknowledgements

We would like to thank Dr. Andrew Kitchen for his time discussing our phylogenetic analyses (trees and tree-thinking), and Dr. Erin Irish for comments on an earlier version of this manuscript.

## Supporting Information

**Figure S1.**
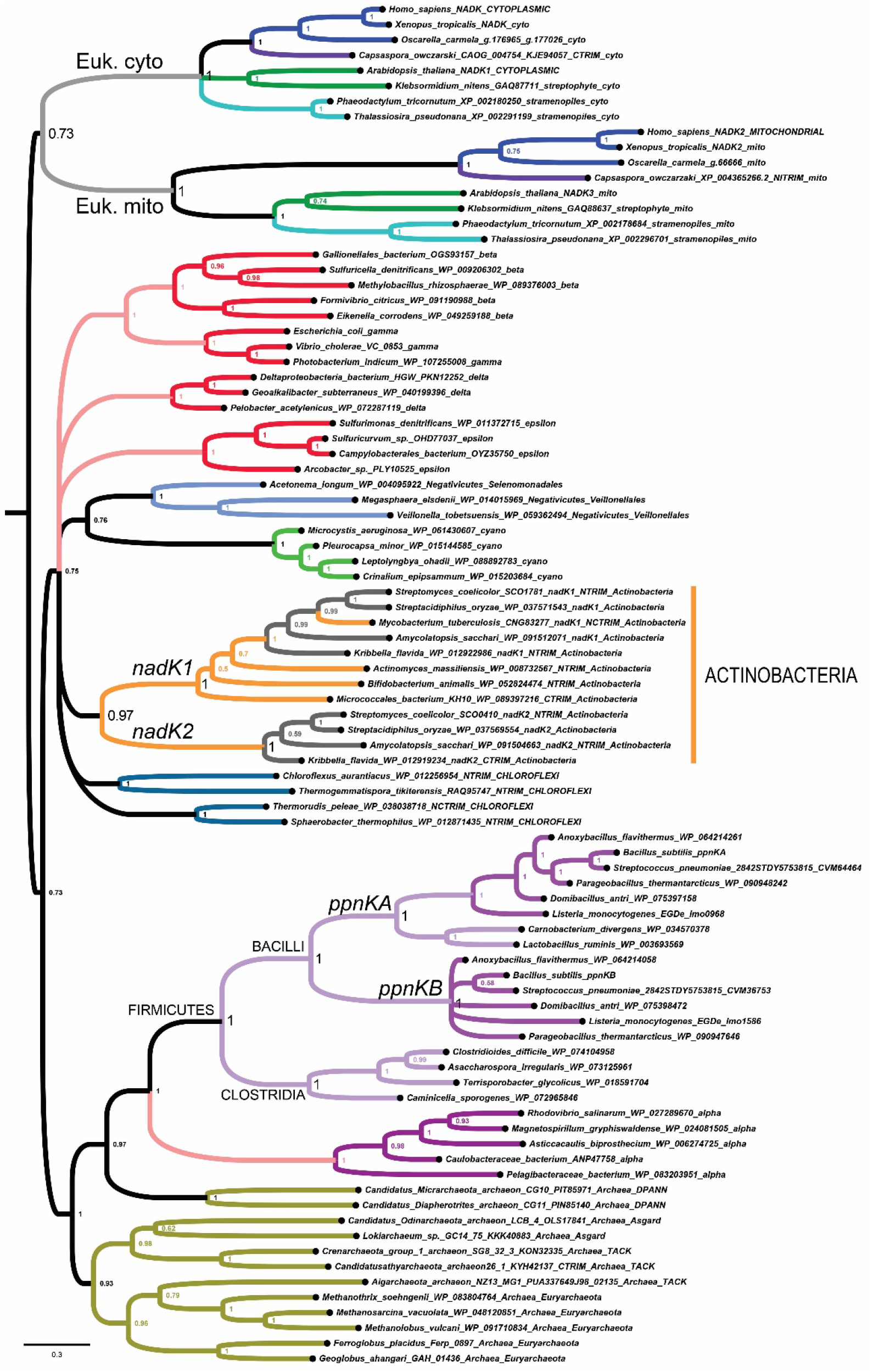
The eukaryotic cyto and mito paralogs are independent duplications from those in the phylum Actinobacteria (orange) and the class Bacilli (phylum Firmicutes, mauve).

**Figure S2.**
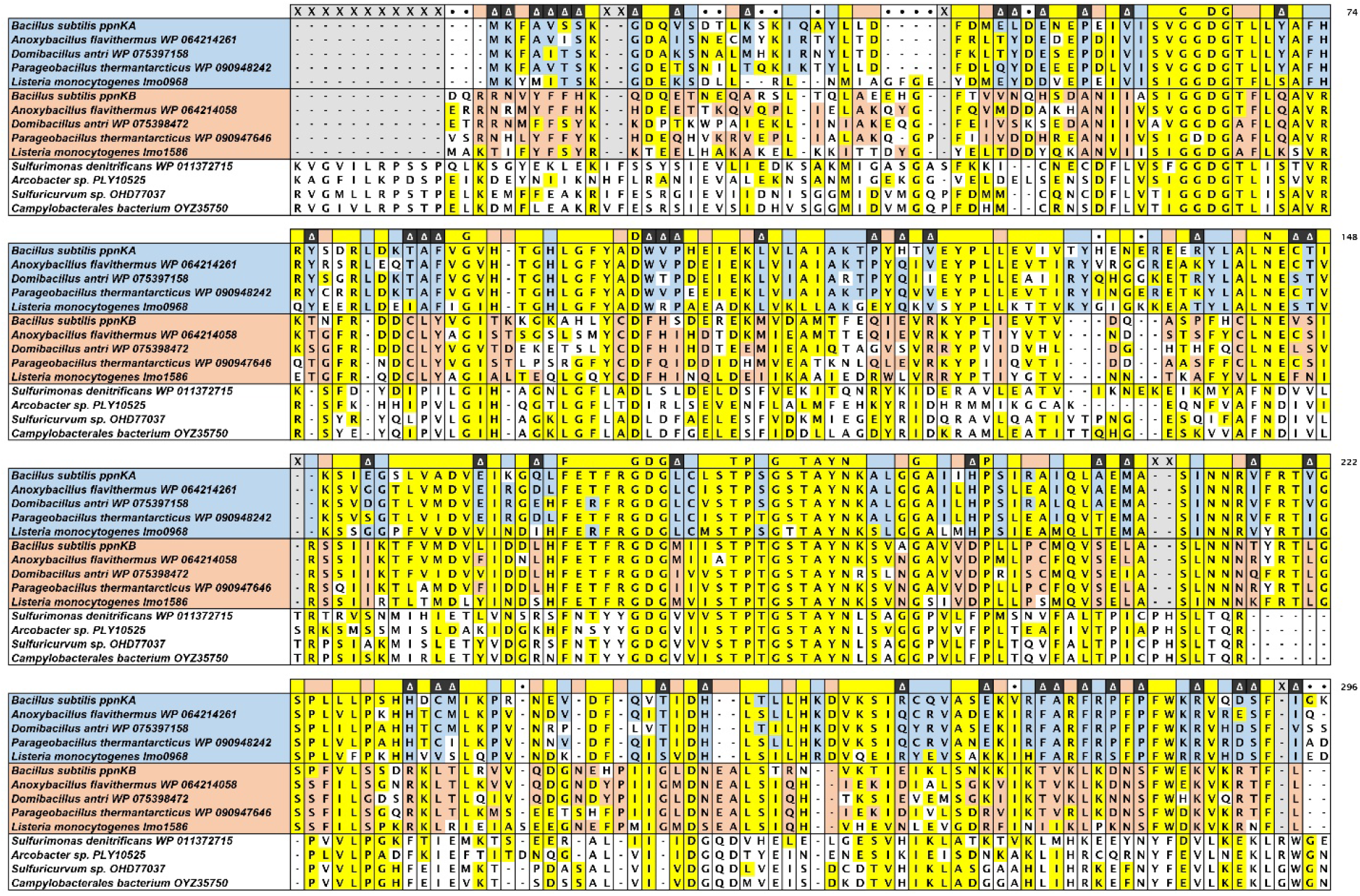
*nadK* paralogs in class Bacilli. The amino acid residues that are differentiated in PpnKA or PpnKB are indicated in blue and red, respectively, while residues that are widely conserved are indicated in yellow.

**Table S1.** Proposed names for *NADK* genes in *Drosophila* and *Caenorhabditis*.

| **Model genetic organism** | **Current systematic gene name** | **Name based on orthology to human** | **Eukaryotic gene clade (sub-clade)** |
| --- | --- | --- | --- |
| *D. melanogaster* | CG6145 | *NADK1a* | “cyto” |
| *D. melanogaster* | CG33156 | *NADK1b* | “cyto” |
| *D. melanogaster* | CG8080 | *NADK2* | “mito” |
| *C. elegans* | Y17G7B.10 | *NADK2a* | “mito” (nematode “nmito”) |
| *C. elegans* | Y77E11A.2 | *NADK2b* | “mito” (nematode “m2c”) |

**Table S2.** List of supplementary files associated with phylogenetic analyses. For each analysis shown, we include the FASTA file (“.fas”), the sequence alignment file (“.masx”), the nexus file (“.nexus”), and associated run files. Each set of files shares the same base file name indicated in the table. Line breaks in FASTA sequences typically indicate places where sequences were trimmed. Some hand curated sequences are documented in WORD files with the base name “CURATION”. All file are included as a compressed archived (“File_S1.tar”).

| **Fig.** | **Base file name** | **Number of**<br>**Metropolis-coupled**<br>**generations** | **Average standard**<br>**deviation of split**<br>**frequencies** |
| --- | --- | --- | --- |
| **1** | NADK_v57 | 800,000 | 0.011 |
| **2** | NADK_v59-fungi-OV | 641,000 | 0.017 |
| **3** | NADK_v63-rando-protists-NCLIP | 1,200,000 | 0.033 |
| **4** | NADK_v58-nematodes | 2,100,000 | 0.031 |
| **5** | NADK_v58-nematodes_only | 1,200,000 | 0.007 |
| **6** | NADK_v62-holozoa (OV) | 1,200,000 | 0.028 |
| **7** | BACILLI_NADK_v108 | 300,000 | 0.009 |
| **S1** | NADK-comparison_v260 | 1,200,000 | 0.018 |

